# Exosomal MFGE8 from high glucose induced endothelial cells is involved in calcification/senescence of vascular smooth muscle cells

**DOI:** 10.1101/2021.09.10.459867

**Authors:** Yu-Qing Ni, Shuang Li, Xiao Lin, Yan-Jiao Wang, Jie-Yu He, Wan-Ling Song, Qun-Yan Xiang, Yan Zhao, Chen Li, Yi Wang, Hua-Hua Li, Zhen Liang, Jun-Kun Zhan, You-Shuo Liu

**Author notes:** **Corresponding Author:** You-Shuo Liu (MD and Ph.D). Department of Geriatrics, Institute of Aging and Age-related Disease Research, The Second Xiangya Hospital, Central South University, 139 Renmin Middle Road, Furong district, Changsha, Hunan, 410011, P.R. China.; Jun-Kun Zhan (MD). Department of Geriatrics, Institute of Aging and Age-related Disease Research, The Second Xiangya Hospital, Central South University, 139 Renmin Middle Road, Furong district, Changsha, Hunan, 410011, P.R. China.

## Abstract

Vascular calcification/aging is a crucial feature of diabetic macro vasculopathy, resulting in serious cardiovascular diseases. The calcification/senescence of vascular smooth muscle cells (VSMCs) induced by hyperglycemia can cause diabetic vascular calcification/aging. However, the mechanism of VSMCs calcification/senescence involved in diabetic vascular calcification/aging remains unknown. The purpose of this study was to determine how the high glucose (HG) information in circulating blood is transmitted from vascular endothelial cells (ECs) to VSMCs, which are not contacted with blood directly. Exosomes have attracted much attention for their vital roles in regulating cell-to-cell communication. In this study, we found that milk fat globule epidermal growth factor 8 (MFGE8) was enriched in high glucose induced human umbilical vein endothelial cell exosomes (HG-HUVEC-Exo) and regulate VSMCs calcification/senescence, characterized by up-regulated expressions of alkaline phosphatase (ALP) and Runt-related transcription factor 2 (Runx2), as well as the increased mineralized nodules and senescence-associated β-galactosidase (SA-β-gal) positive cells. Upstream mechanism studies showed that sirtuin1 (SIRT1) was involved in VSMCs calcification/senescence by affecting the expression of MFGE8. We also found that inflammatory response mediated by IL-1β, IL-6, and IL-8 was closely associated with MFGE8 and played a key role in regulating HG-HUVEC-Exo-induced VSMCs calcification/senescence. These findings provide a new insight into the mechanism of exosomal MFGE8 as a potential preventive and therapeutic target for the intervention of diabetic vascular calcification/aging.

## Introduction

Diabetes and vascular-related complications are major causes of disability and death of patients that threaten human public health. However, there is no evidence of long-term benefits of intensive blood glucose control on macrovascular events and mortality (Zoungas et al., 2014), suggesting that the prevention and treatment of diabetic vascular diseases is more than just glucose control. Vascular aging is a special organic aging, whereas vascular calcification is an important phenotype of vascular aging. Vascular calcification/aging is a crucial cause of diabetic vascular-related complications (Currie and Delles, 2017). Therefore, it is of great significance to study the mechanism of vascular calcification/aging in diabetes.

Alterations in the biological functions and structural properties of endothelial cells (ECs) and vascular smooth muscle cells (VSMCs) are the cellular basis of vascular calcification/aging. Cellular senescence is characterized by a series of physiological and morphological changes, which can be evaluated by senescence-associated β-galactosidase (SA-β-gal) staining and the expression of aging-related proteins, such as p16, p53 and p21 (Hernandez-Segura et al., 2018). Osteogenic hallmarks including alkaline phosphatase (ALP) and Runt-related transcription factor 2 (Runx2), as well as Alizarin Red S staining can be used to detect cellular calcification (Chen et al., 2020). VSMCs calcification/senescence in the media of vessel wall is the main manifestation of diabetic vascular calcification/aging (Lin et al., 2019b). However, how is high glucose (HG) information transmitted from ECs to VSMCs that are not contacted with circulating blood directly?

Exosomes, newly identified natural nanocarriers (30-150 nm) and intercellular messenger, have attracted much attention for their vital role in regulating cell-to-cell communication (Xiong et al., 2021; Yáñez-Mó et al., 2015). Because of the variety and abundance of specific cargos, such as proteins, lipids, nucleic acids, exosomes are involved in diverse physiological and pathophysiological processes including proliferation, migration, differentiation, homeostasis, apoptosis, and senescence (Jenjaroenpun et al., 2013; Liao et al., 2013). The active interaction and crosstalk between ECs and VSMCs is the key to modulate vascular aging process, which is achieved through the release of exosomes from ECs (Ni et al., 2020a; Zhao et al., 2017). Moreover, our previous studies indicated that HG-induced human umbilical vein endothelial cell exosomes (HG-HUVEC-Exo) stimulated VSMCs calcification/senescence (Li et al., 2019; Lin et al., 2019a). However, the overall mechanism of the interaction between ECs-derived exosomes and target VSMCs remains largely uncharacterized. Which cargo in exosomes is involved in the communication between ECs and VSMCs in hyperglycemia, and what is the signaling pathway? It is of great significance to further study the roles and mechanisms of HG-HUVEC-Exo in regulating VSMCs calcification/senescence.

In this study, we identified that exosomal milk fat globule epidermal growth factor 8 (MFGE8) could be carried and integrated to VSMCs, thereby modulating VSMCs calcification/senescence. Further mechanism studies revealed that sirtuin1 (SIRT1) is down-regulated in HUVEC-Exo under hyperglycemia, leading to the up-regulation of exosomal MFGE8, which in turn influence VSMCs calcification/senescence through promoting inflammatory response mediated by IL-1β, IL-6, and IL-8. Taken together, our findings provide a potential therapeutic target for diabetic vascular calcification/aging.

## Results

### The indirect effects of HG-HUVEC-Exo on VSMCs calcification/senescence

Firstly, we constructed a HG-HUVEC induced calcification/senescence model of VSMCs. VSMCs were co-cultured with NG (5 mmol/l) and HG (30 mmol/l)-HUVEC for 0, 24 or 48h. Compared with VSMCs treated with NG-HUVEC, the expressions of both ALP and Runx2 were greatly increased in VSMCs treated with HG-HUVEC for 48h (Fig. 1A). Furthermore, the mineralized nodules (Fig. 1B) and senescent cells (Fig. 1C) in VSMCs determined via Alizarin Red S staining and SA-β-gal staining were also significantly increased in HG-HUVEC induced VSMCs for 14 days. We hypothesized that exosomes were associated with the communication of HUVEC and VSMCs. To indirectly demonstrate the effects of HUVEC-Exo in regulating VSMCs calcification/senescence under hyperglycemia, HG-HUVEC were pre-treated with GW4869 to inhibit the secretion of exosomes. The results showed that GW4869 pre-treated HG-HUVEC had little effect on VSMCs calcification/senescence (Fig. 1A-C). These results consequential suggested that exosomes, rather than other contents secreted by HUVEC, promoted VSMCs calcification/senescence under hyperglycemic condition.

**Figure 1.**
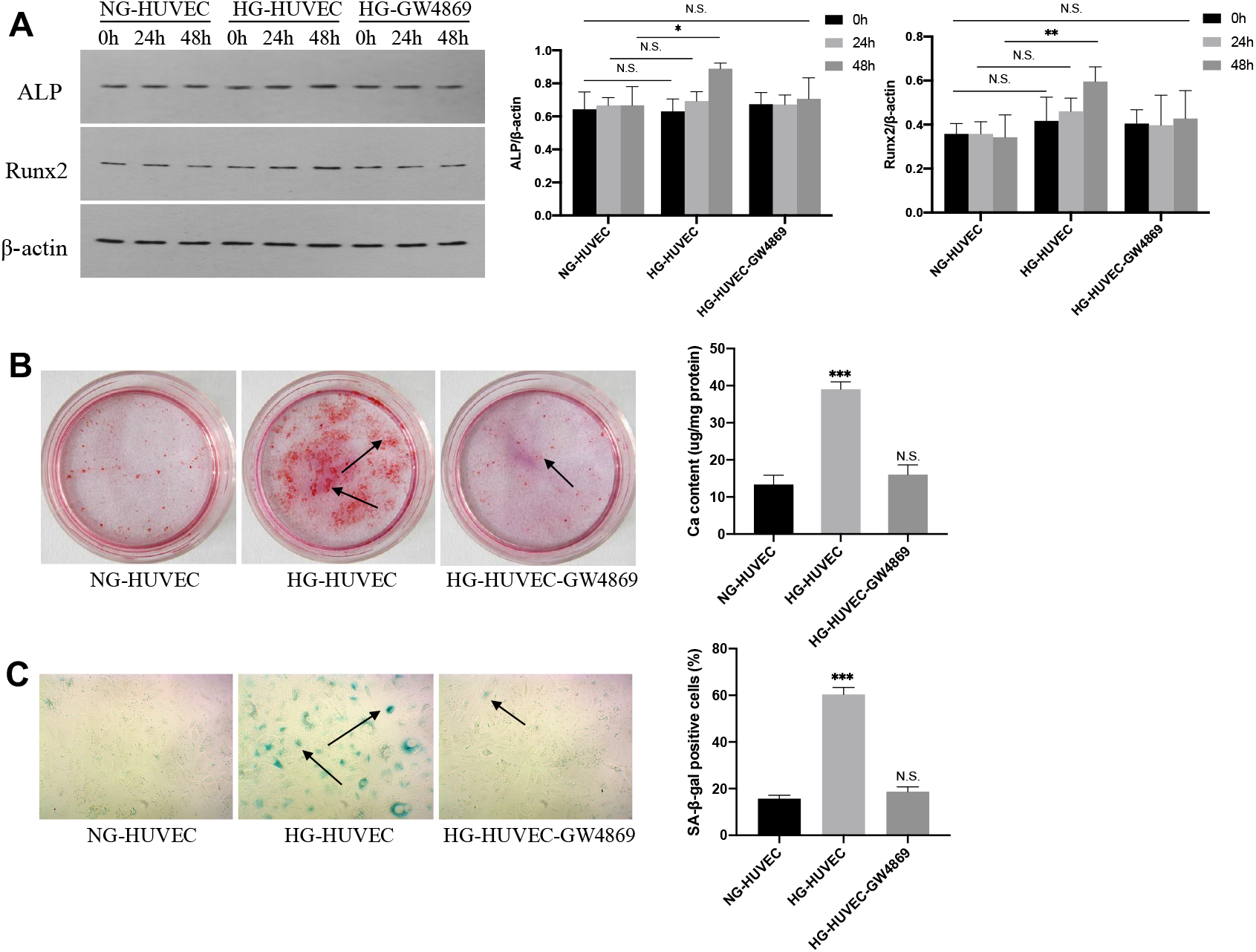
HG-HUVEC stimulate VSMCs calcification/senescence. (A) The expression of ALP, Runx2 were measured by western blot analysis in VSMCs under conditions of NG, HG, or HG with GW4869 for 0h, 24h, 48h. (B) Alizarin Red S staining and (C) SA-β-gal staining showed the calcium content and senescent cells in VSMCs. In panel B, the content of calcium was extracted by cetylpyridinium chloride and quantified by spectrophotometry; the arrow represented calcium nodules. Semiquantitative analysis of SA-β-gal staining positive cells was performed by ImageJ in panel C; the arrow showed the senescent VSMCs. The data are expressed as the mean ± SD (n = 3, ^*^*P* < 0.05, ^**^*P* < 0.01, ^***^*P* < 0.001; N.S., no significance; NG, normal glucose; HG, high glucose).

### HG-HUVEC-Exo induce VSMCs calcification/senescence

To further assess the effects of exosomes on VSMCs calcification/senescence, we isolated exosomes from the supernatants of HUVEC treated with NG and HG to directly interfere with VSMCs. Transmission electron microscope showed the bilayer structure of the vesicles (Fig. 2A), and nanoparticle tracking analyzer detected a mean diameter of 133.64 ± 5.24 nm of vesicles (mean ± SEM) (Fig. 2B). Western blot identified the expression of exosomal-specific markers CD63 and CD9 on the exosome surface as well (Fig. 2C). These characteristics are accordance with previous reports on exosomes (Xiong et al., 2021), confirming that the vesicles were exosomes. After extraction and identification of exosomes, we further exploited the ability of HUVEC-Exo to regulate VSMCs calcification/senescence in hyperglycemia. VSMCs were co-cultured with the presence of HUVEC-Exo. As shown in (Fig. 2D-F), HG-HUVEC-Exo aggravated calcification/senescence of VSMCs, which was validated by significant increase in the protein levels of ALP and Runx2, and the increase in formations of mineralized nodule and SA–β-gal positive cells.

**Figure 2.**
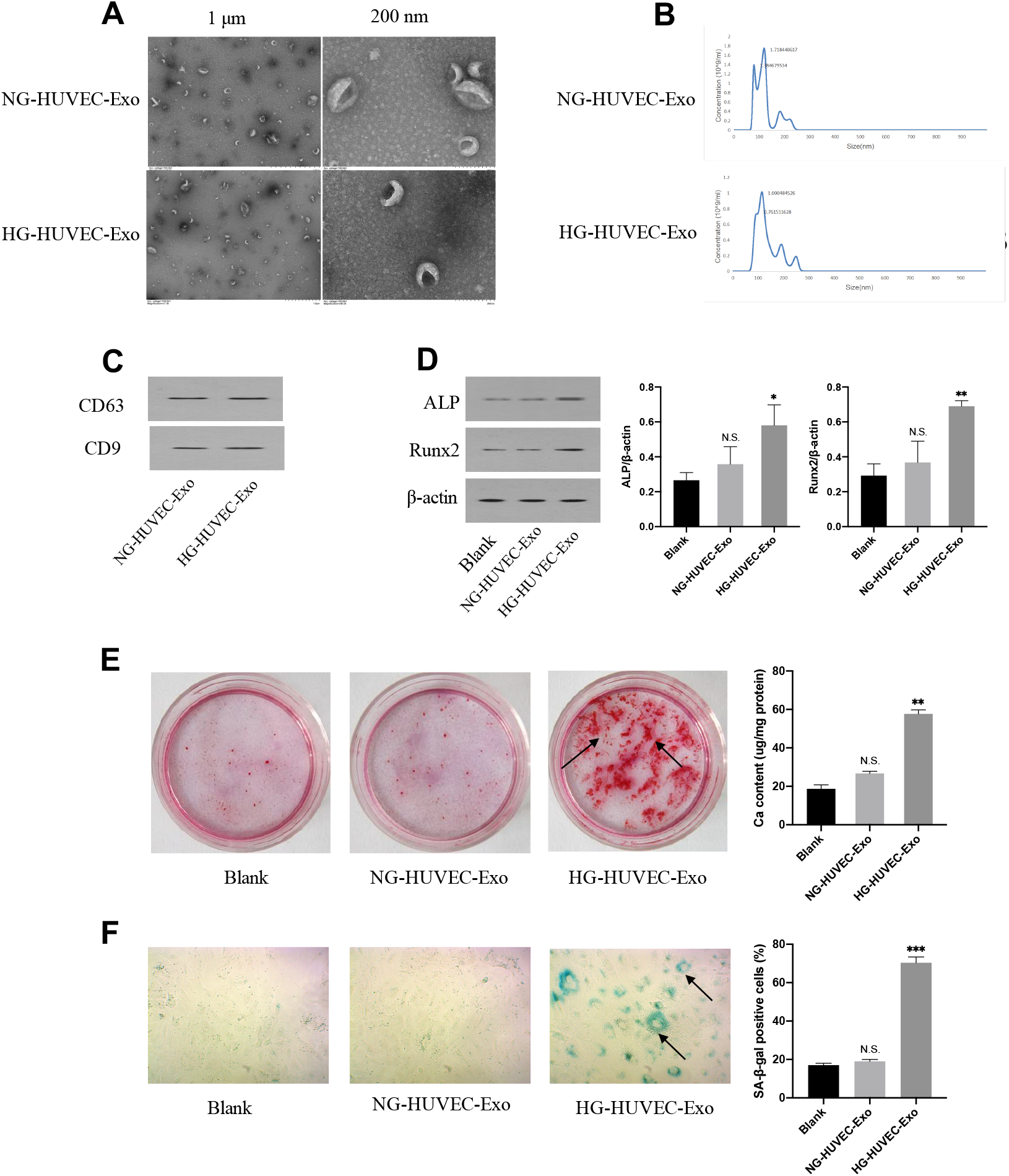
The effects of HG-HUVEC-Exo on VSMCs calcification/senescence. Transmission electron microscope, nanoparticle tracking analyzer, and western blot were applied in identification of exosomes. (A) Transmission electron microscope was used to assess the morphology of exosomes treated with NG and HG. Scar bar represents 1 μm and 200 nm, respectively. (B) The size and concentration of exosomes were measured by nanoparticle tracking analyzer. (C) Exosomal specific marker CD63 and CD9 were detected by western blot. (D) The protein expression levels of ALP and Runx2 were assayed by western blot in VSMCs. (E and F) Calcification and senescence of VSMCs were examined via Alizarin Red S staining and SA-β-gal staining. The data are expressed as the mean ± SD (n = 3, ^*^*P* < 0.05, ^**^*P* < 0.01, ^***^*P* < 0.001; N.S., no significance).

### MFGE8 is enriched in HG-HUVEC-Exo

HUVEC-Exo were involved in the modulation of VSMCs calcification/senescence. Therefore, it was then decided to investigate the roles of molecule in HG-HUVEC-Exo on VSMCs function. Proteomics was used to analyze differential protein expression profiles between HG-HUVEC-Exo and NG-HUVEC-Exo. A total of 27 up-regulated and 68 down-regulated heteroproteins were identified using a corrected *P* value < 0.05. Then, intersections of up-regulated proteins in HG-HUVEC-Exo in proteomics were analyzed with Disease Database (https://rgd.mcw.edu/wg/portals/) of diabetes mellitus and vascular disease, 5 intersection genes were screened out (Fig. 3A and B). Finally, the target protein was preliminarily determined as MFGE8 through literature review in PubMed (https://pubmed.ncbi.nlm.nih.gov/). Further western blot confirmed the enrichment of MFGE8 in HG-HUVEC-Exo compared with those treated with NG-HUVEC-Exo (Fig. 3C).

**Figure 3.**
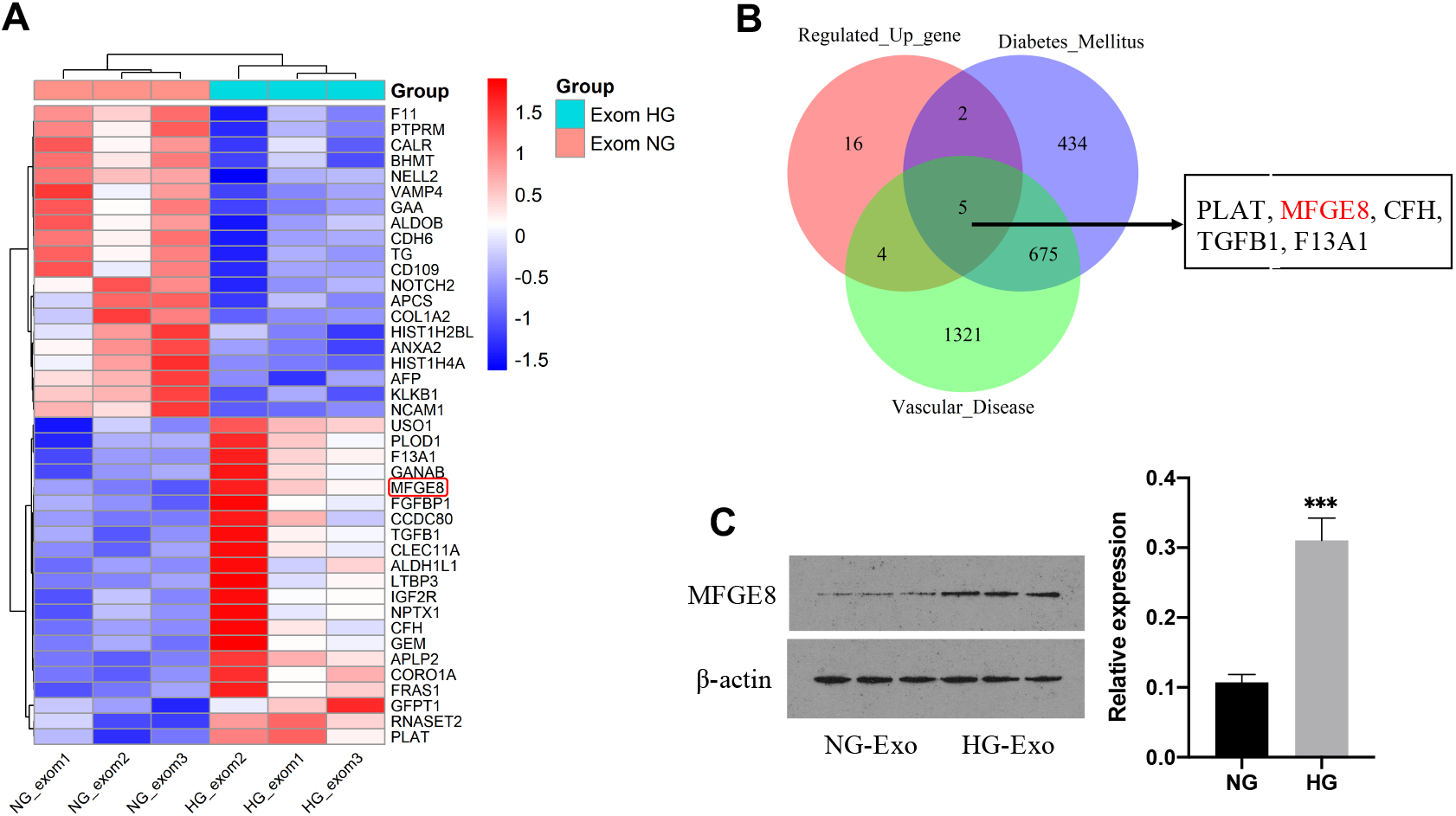
Screening and validation of exosomal MFGE8. (A) Proteomics was used to analyze the differentially expressed proteins between NG-HUVEC-Exo and HG-HUVEC-Exo (a corrected *P* value < 0.05). (B) Bioinformatics analysis was carried out to screen the intersected differentially expressed genes. (C) Western blot was conducted to verify the differential expression of MFGE8 in NG-HUVEC-Exo and HG-HUVEC-Exo. The data are expressed as the mean ± SD (n = 3, ^***^*P* < 0.001).

### HUVEC-Exo carry MFGE8 to induce VSMCs calcification/senescence

Next, fluorescent microscopy was used to verify whether HUVEC-Exo could carry MFGE8 to VSMCs. DiD-labelled exosomes showed red spotty staining while GFP-labelled MFGE8 represented green spotty staining. The nucleus was counterstained as blue dots. Merged fluorescence appeared orange in VSMCs. It indicated that DiD-labelled HUVEC-Exo could carry GFP-labelled MFGE8 to VSMCs (Fig. 4A). An overexpression vector (OV) of MFGE8 (MFGE8-OV) or an empty negative control (NC) of MFGE8 (MFGE8-NC) was transfected into HUVEC. Exosomes derived from HUVEC were isolated and co-cultured with VSMCs. Western blot analysis showed that the relative expression of MFGE8 in VSMCs treated with Exo-MFGE8-OV was four-fold than that in the Exo-MFGE8-NC group (Fig. 4B), which further confirmed that HUVEC derived exosomal MFGE8 could be absorbed and integrated into VSMCs. To determine the role of exosomal MFGE8 in VSMCs calcification/senescence, we verified the direct and indirect effects of HUVEC-MFGE8 and HUVEC-Exo on VSMCs, respectively. First, HUVEC were transfected with MFGE8-OV or MFGE8-NC plasmids for 48h and then transfected with VSMCs in co-culture cell chamber. VSMCs were collected and detected for various indicators, including ALP, Runx2, Alizarin Red S staining and SA-β-gal staining. The results showed that MFGE8 were able to promote VSMCs calcification/senescence, characterized by up-regulated expressions of ALP and Runx2, as well as the increased mineralized nodules and SA-β-gal positive cells (Fig. 4C-E). In addition, compared with those treated with NG-HUVEC-Exo, the levels of ALP, Runx2, mineralized nodules and SA-β-gal positive cells were significantly increased in VSMCs treated with HG-HUVEC-Exo. As expected, the roles of HG-HUVEC-Exo on VSMCs calcification/senescence were markedly attenuated when HG-HUVEC-Exo knocked down MFGE8 using SiRNA-MFGE8, which was confirmed by decreases in ALP, Runx2, Alizarin Red S staining and SA–β-gal staining (Fig. 4F-H). Altogether, these data indicated that exosomal MFGE8 played a pivotal role in VSMCs calcification/senescence.

**Figure 4.**
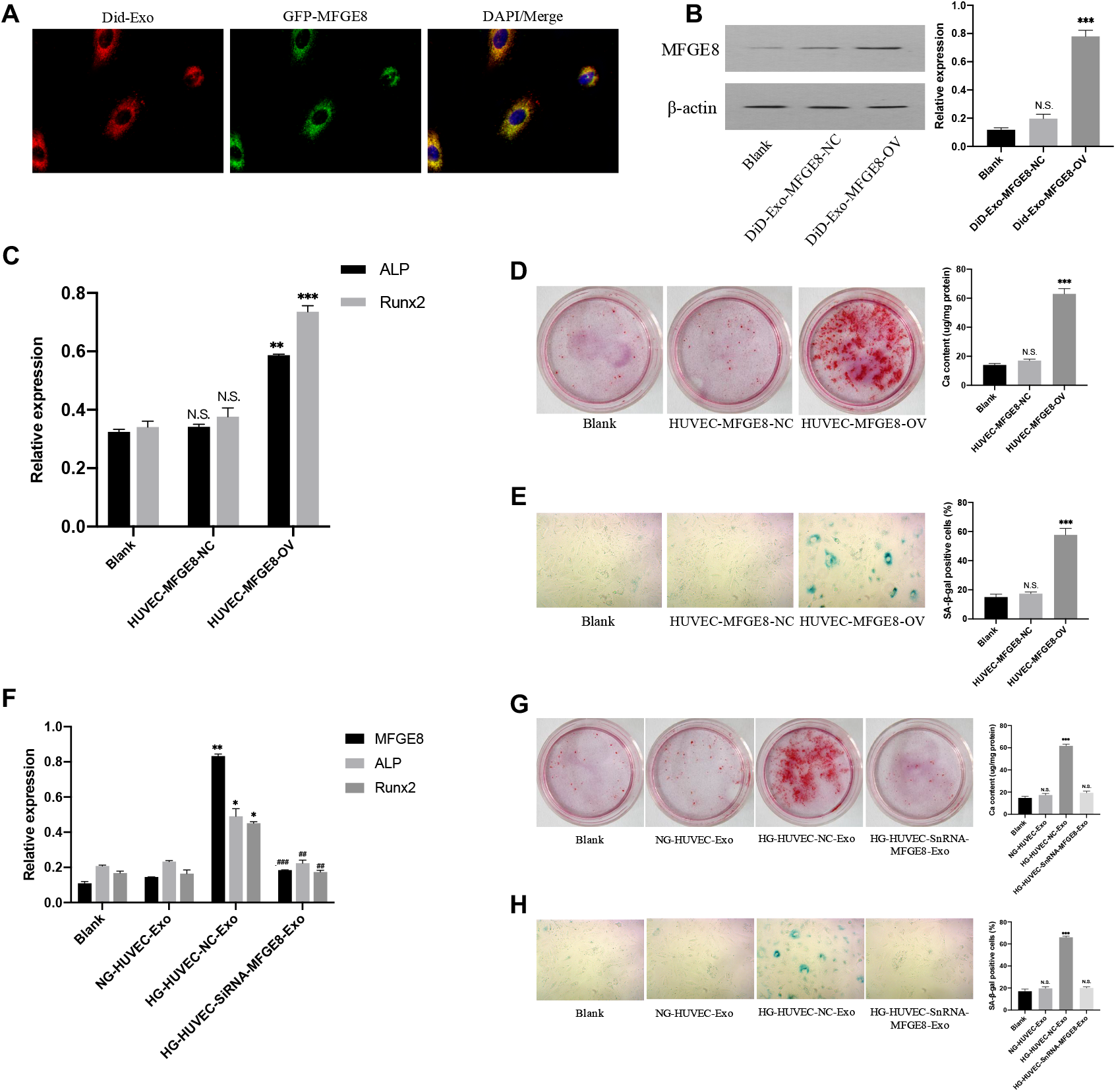
Exosomal MFGE8 is involved in regulating VSMCs calcification/senescence. (A) Fluorescent microscopy revealed that DiD-labelled exosomes could carry GFP-labelled MFGE8 to VSMCs. (B) Western blot verified that the expression of MFGE8 was increased under the condition of MFGE8-OV. (C) The protein expression levels of ALP and Runx2 were assayed by western blot. VSMCs were co-cultured with HUVEC-MFGE8-NC or HUVEC-MFGE8-OV for 48h. (D) Alizarin Red S staining showed the calcification in VSMCs. (E) SA-β-gal staining was used to determine senescence VSMCs. (F) VSMCs were co-cultured with NG-HUVEC-Exo, HG-HUVEC-NC-Exo or HG-HUVEC-SiRNA-MFGE8-Exo, and the expression of MFGE8, ALP, Runx2 were measured by western blot analysis. (G and H) Alizarin Red Staining and SA-β-gal staining were used to determine the calcification and senescence of VSMCs. The data are expressed as the mean ± SD (n = 3, compared with NG-HUVEC-Exo, ^*^*P* < 0.05, ^**^*P* < 0.01, ^***^*P* < 0.001; N.S., no significance; compared with HG-HUVEC-NC-Exo, ^##^*P* < 0.01; ^###^*P* < 0.001; NC, negative control; OV, overexpression vector).

### SIRT1 regulates MFGE8 and influences VSMCs calcification/senescence

Why is the expression of MFGE8 elevated in HG-HUVEC-Exo and what is its regulatory mechanism? In order to explore the upstream mechanism, RGD disease database analysis (https://rgd.mcw.edu/wg/portals/) was utilized. It was found that there were 22 intersection genes of diabetes mellitus, vascular disease, hyperglycemia, and diabetic angiopathies. Sirt1 aroused our great interest, not only because it is a “star molecule” in aging, but it is also the only protease among 22 intersection genes (Fig. 5A). The mRNA expression of MFGE8 was significantly increased under the condition of HG, whereas that of SIRT1 was decreased by qRT-PCR (Fig. 5B and C). SRT2104, an activator of SIRT1, was utilized to promote the activity of SIRT1 in HUVEC. Interestingly, the results showed that both the mRNA and protein expression of MFGE8 were decreased in HG-HUVEC treated with SRT2104 (Fig. 5D and E), suggesting a potential regulatory role of SIRT1 on MFGE8. To further clarify the correlation between SIRT1 and MFGE8, we then performed ChIP-PCR assay. The results revealed that the enrichment of MFGE8 sequence was markedly increased when using SIRT1 antibodies (including H3K9Ac or H3K27Ac) for immunoprecipitation compared with a control IgG antibody (Fig. 5F). Moreover, compared with HG-HUVEC, the enrichment of MFGE8 sequences with SIRT1 antibodies were decreased in HG-HUVEC treated with SRT2104 (Fig. 5F). Next, we further detected the role of SIRT1 in VSMCs calcification/senescence. The expression of MFGE8 in VSMCs decreased or increased after the addition of SIRT1 agonist (SRT2104) or inhibitor (Selisistat), respectively (Fig. 5G). Besides, compared with control group, HUVEC treated with SRT2104 attenuated VSMCs calcification/senescence, while HUVEC treated with Selisistat accelerated VSMCs calcification/senescence, which were verified by the significant changes in the expression of ALP, Runx2, Alizarin Red S staining and SA– β-gal staining (Fig. 5G-I). Taken together, these data suggested that SIRT1 could regulate VSMCs calcification/senescence by affecting the expression of MFGE8.

**Figure 5.**
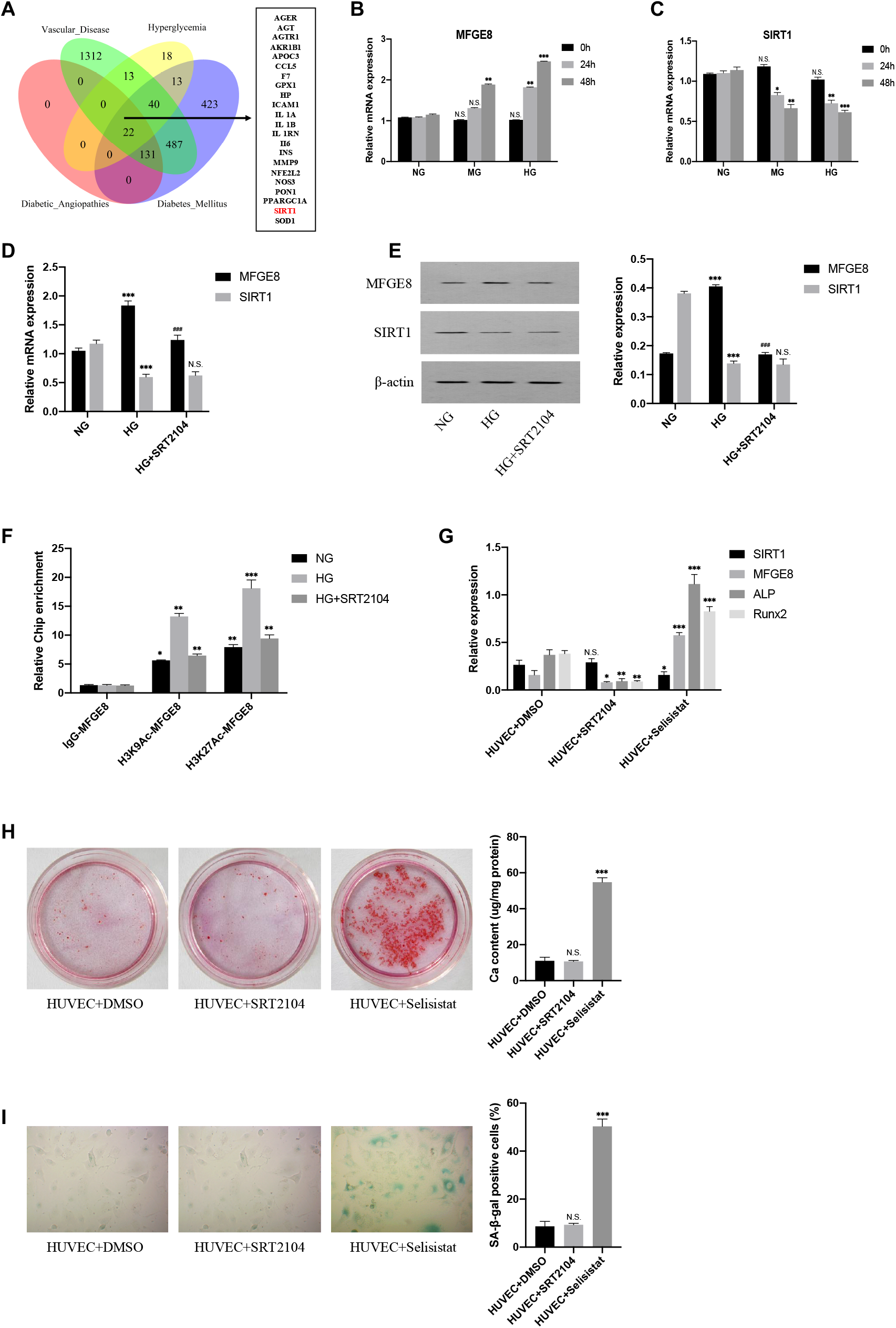
SIRT1 regulates MFGE8 and influences the calcification/senescence of VSMCs. (A) The RGD disease database was utilized to analyze the intersection genes of diabetes mellitus, vascular disease, hyperglycemia, and diabetic angiopathies. (B) MFGE8 and (C) SIRT1 mRNA expression were assessed by qRT-PCR. (D and E) QRT-PCR and immunoblotting were used to detect mRNA and protein levels of MFGE8 and Sirt1 in HUVEC. (F) ChIP assay using H3K9Ac, H3K27Ac or normal IgG antibodies and PCR performed with primers for promoter region of MFGE8 gene were used to verify the correlation between SIRT1 and MFGE8. (G) Western blot was used to detect the expression of SIRT1, MFGE8, ALP and Runx2 in VSMCs co-cultured with HUVEC pre-treated by DMSO, SRT2104, Selisistat, respectively. (H and I) Alizarin Red S staining and SA-β-gal staining showed the calcium content and senescent cells in VSMCs under different experimental interventions. The data are expressed as the mean ± SD (n = 3, ^*^*P* < 0.05, ^**^*P* < 0.01, ^***^*P* < 0.001; N.S., no significance).

### Inflammatory response plays a key role in HG-HUVEC-Exo-MFGE8 induced VSMCs calcification/senescence

To explore the downstream signaling pathway of MFGE8 in HG-HUVEC-Exo to promote VSMCs calcification/senescence, we conducted bioinformatics analysis on the differentially expressed genes of VSMCs treated with HG-HUVEC-Exo and NG-HUVEC-Exo. The volcano plot visually showed the differential gene expressions between the two groups, among which the levels of IL-1β, IL-6, IL-8, and MFGE 8 were all up-regulated in HG treatment group (Fig. 6A). Moreover, the bubble plot of *GO* enrichment analysis showed an enrichment of up-regulated differential expression genes in the inflammatory response (Fig. 6B). In addition, the expression levels of IL-1β, IL-6 and IL-8 in HG-HUVEC-Exo were up-regulated by ELISA. Interestingly, downregulation of MFGE8 by siRNA resulted in a significant decrease in IL levels (Fig. 6C). Collectively, these findings suggest that exosomal MFGE8 derived from HG-induced HUVEC on VSMCs calcification/senescence through an inflammatory signaling pathway mediated by IL-1β, IL-6 and IL-8.

**Figure 6.**
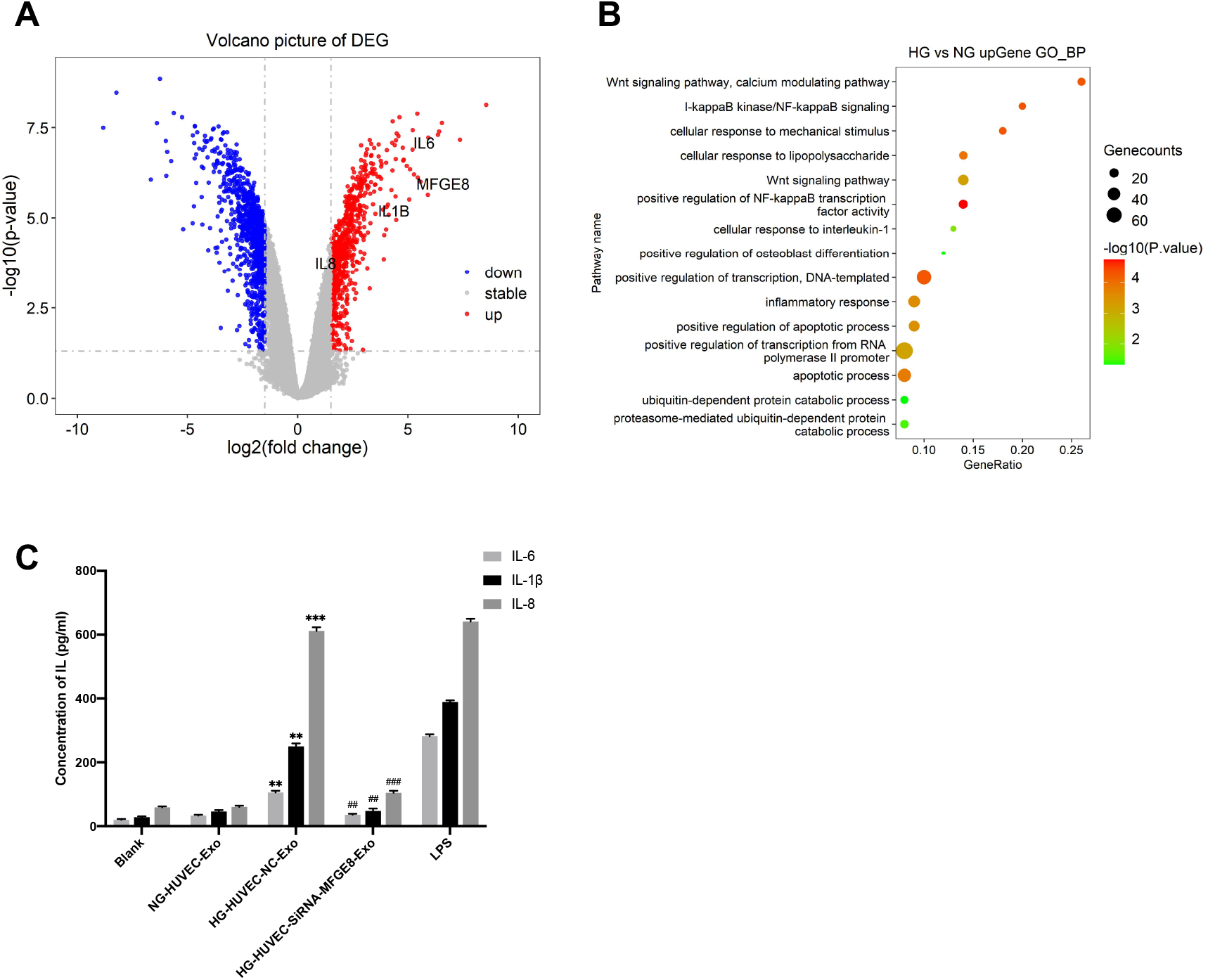
Inflammatory factors are involved in HUVEC-Exo-MFGE8 induced VSMCs calcification/senescence. (A) Volcano plot depicted the differentially expressed genes in VSMCs treated with HG-HUVEC-Exo and NG-HUVEC-Exo. Each dot represented a gene, where red and blue dots denoted genes with high or low expression levels, respectively. (B) Bubble plot of *GO* enrichment analysis was conducted to identify the biological processes involving differential gene expression. It showed an enrichment bubble plot of up-regulated differential expression genes, where the size of the dots represented the number of genes enriched in the pathway, and the color of the dots described the degree of enrichment. (C) ELISA was used to determine the differential expression levels of IL-1β, IL-6 and IL-8 from the supernatants of VSMCs treated with NG-HUVEC-Exo, HG-HUVEC-NC-Exo, HG-HUVEC-SnRNA-MFGE8-Exo and LPS.

## Discussion

The main hazard of diabetes is its vascular-related complications, which is the major causes of disability and death in patients. Vascular calcification/aging has been identified as a serious risk factor for vascular complications, which can be significantly aggravated by hyperglycemia (Stabley and Towler, 2017). Vascular aging is characterized by morphological abnormalities such as cell enlargement and vacuole formation at the cellular level. Histologically and functionally, it is manifested as disordered elastic fibers, increased collagen fibers, widened pulse pressure, and elevated pulse wave velocity (Society, 2018). Vascular calcification, also termed as Monckeberg’s calcification, is an pivotal phenotype of vascular aging, mainly involving the VSMCs in the media of vessels (Lin et al., 2018; Ryu et al., 2021). Since ECs and VSMCs are important components of the inner and middle layers of blood vessel, the changes in structure and function of ECs and VSMCs are closely related to vascular dysfunction. It has been shown that VSMCs calcification/senescence exhibits a key role in the occurrence and development of diabetic vascular calcification/aging (Zhao et al., 2014). Our research group has been intensively engaged in this field and has made a series of original achievements (Li et al., 2019; Lin et al., 2019a; Lin et al., 2019b; Tan et al., 2016; Zhan et al., 2014; Zhan et al., 2018; Zhan et al., 2015). However, how is communicating information transmitted from ECs to VSMCs that are not in direct contact with circulating blood?

Recently, it has been reported that exosomes play a vital role in regulating vascular functions and pathologies via achieving the close interaction between ECs and VSMCs (He et al., 2018). We found that exosomes secreted by melatonin-treated VSMCs alleviated the senescence and osteogenic differentiation of VSMCs (Xu et al., 2020). Emerging evidence has revealed that ECs-derived exosomes regulate VSMCs phenotypic modulation, including cellular migration, calcification, apoptosis, and senescence (Chen et al., 2013; Maegdefessel et al., 2015). For instance, Shan *et al*. demonstrated that exosomal RNCR3 produced by ox-LDL treated ECs mediated VSMCs proliferation and migration, and regulated VSMCs function, leading to atherosclerotic phenotype (Shan et al., 2016). In this study, we found that HG-HUVEC aggravated VSMCs calcification/senescence, which was confirmed by significant increase in the protein levels of ALP and Runx2, as well as the increase in formations of Alizarin Red S staining and SA-β-gal staining in VSMCs. Meanwhile, inhibition of exosomes by GW4869 partially alleviated this induction, suggesting the importance of HUVEC-Exo in the regulation of VSMCs calcification/senescence under hyperglycemia. Furthermore, we isolated and identified exosomes using morphology, size and specific markers. We verified that HUVEC-Exo were involved in the regulation of HG-induced aging-related phenotype and calcification of VSMCs. Therefore, exosomes could mediate HG-induced VSMCs calcification/senescence and serve as an effective bioinformation transporter.

MFGE8, a multifunctional glycoprotein, has attract much attention over the last few years. It has been identified as a mediator in cell-to-cell interactions involved in diverse processes, including tumorigenesis, angiogenesis, and innate immune response (Yang et al., 2011). Additionally, high-throughput proteomic screening and cellular network analysis have determined MFGE8 as the cause of aging-related increases in extracellular matrix adhesion molecules (Wang et al., 2014). Although recent studies have found that MFGE8 is related to aging, there are only a few reports on its relationship with VSMCs. Recently, we first demonstrated the roles and mechanisms of MFGE8 in vascular aging-related diseases with special emphasis on the functions of ECs and VSMCs (Ni et al., 2020b). However, the association between MFGE8 and diabetic vascular calcification/aging has not been reported to date. In this study, proteomics analysis showed that MFGE8 was enriched in HG-HUVEC-Exo compared with NG-HUVEC-Exo, which was confirmed by western blot analysis. To verify that HUVEC-Exo could carry MFGE8 to VSMCs, DiD and GFP were used to label exosomes and MFGE8, respectively. Interestingly, fluorescent microscopy showed that these HUVEC-derived exosomal MFGE8 were internalized by VSMCs. Of note, the relative expression of MFGE8 was significantly increased in VSMCs treated with Exo-MFGE8-OV, further demonstrating that HUVEC-derived exosomal MFGE8 could be absorbed and integrated into VSMCs. Moreover, we found that exosomal MFGE8 promoted the expressions of ALP and Runx2, as well as the increased mineralized nodules and SA-β-gal positive cells under hyperglycemia. The inductions were almost eliminated by siRNA knockout of MFGE8 expression in HUVEC. These results suggest that HG-HUVEC-Exo induce the calcification and senescence of VSMCs by carrying MFGE8.

Sirtuins are a family of nicotinamide adenine dinucleotide (NAD+)-dependent deacetylases that are involved in a variety of cellular processes (Singh et al., 2018). SIRT1 is a “star molecule” in aging and plays key roles in mitochondrial function, DNA repair, metabolism and antioxidant activity (Hall et al., 2013; Morigi et al., 2018). It functions through the deacetylation of histone proteins H1, H3 and H4, as well as non-histone targets such as nuclear factor-kappa B (NF-kB) and p53 (Nakagawa and Guarente, 2014). Here, RGD disease database analysis found that Sirt1 is associated with diabetes mellitus, vascular disease, hyperglycemia, and diabetic angiopathies. However, the correlation between SIRT1 and MFGE8 during aging remains unclear. In the current study, we found that the mRNA expression of MFGE8 and SIRT1 were inversely parallel in HG-HUVEC, and MFGE8 level could be regulated by SIRT1 agonist or inhibitor. Furthermore, ChIP-PCR assay showed that the enrichment of MFGE8 sequence was significantly increased with SIRT1 antibody immunoprecipitation, whereas it was markedly decreased in SRT2104 treated group. We also examined the role of SIRT1 in VSMCs calcification/senescence using SRT2104 and Selisistat. Taken together, these data demonstrates that SIRT1 regulate VSMCs calcification/senescence by affecting the expression of MFGE8. However, the specific signalling pathway of HG-HUVEC-derived exosomal MFGE8 to promote the calcification/senescence of VSMCs remains to be further studied.

It is well known that inflammatory response and inflammatory cytokines are involved in vascular calcification/aging, and vascular calcification/aging can further aggravate inflammation, thereby forming a vicious cycle (Ungvari et al., 2018; Wang et al., 2014). Increasing evidences suggest that proinflammatory cytokines such as IL-1β, IL-6, IL-8, contribute to the promotion of VSMCs calcification, leading to vascular calcification (Bouabdallah et al., 2019; Lee et al., 2019). Khalifeh et al. proved that MFGE8 was involved in VSMCs senescence by regulating the expression of inflammatory cytokines (Khalifeh-Soltani et al., 2018). In this regard, bioinformatics analysis was performed on the differentially expressed genes in VSMCs treated with HG-HUVEC-Exo and NG-HUVEC-Exo. We found that MFGE8, IL-1β, IL-6, IL-8 were all highly expressed. GO analysis mainly includes three categories including biological process, molecular function, and cell component. Bubble plot showed that the inflammatory response was one of the important signal transduction pathways. Moreover, we further found that the expression levels of IL-1β, IL-6 and IL-8 were up-regulated in HG-HUVEC-Exo. Downregulation of MFGE8 resulted in a significant decrease in IL levels in VSMCs. Collectively, these findings suggest that the inflammatory response mediated by IL-1β, IL-6 and IL-8 may be one of the regulatory mechanisms of exosomal MFGE8 derived from HG-induced HUVEC on the calcification/senescence of VSMCs.

In summary, our findings revealed a vital role of exosomal MFGE8 in regulating diabetic vascular calcification/aging. Exosomal MFGE8 from HG-HUVEC was regulated by SIRT1 and influenced VSMCs calcification/senescence. In addition, HG-HUVEC derived exosomal MFGE8 promoted VSMCs calcification/senescence through inflammatory signalling pathway mediated by IL-1β, IL-6, and IL-8 (Fig. 7). These interesting findings provide future directions for the development of novel interventions to prevent diabetic vascular calcification/aging by targeting exosomal MFGE8 in fundamental molecular and cellular aging processes.

**Figure 7.**
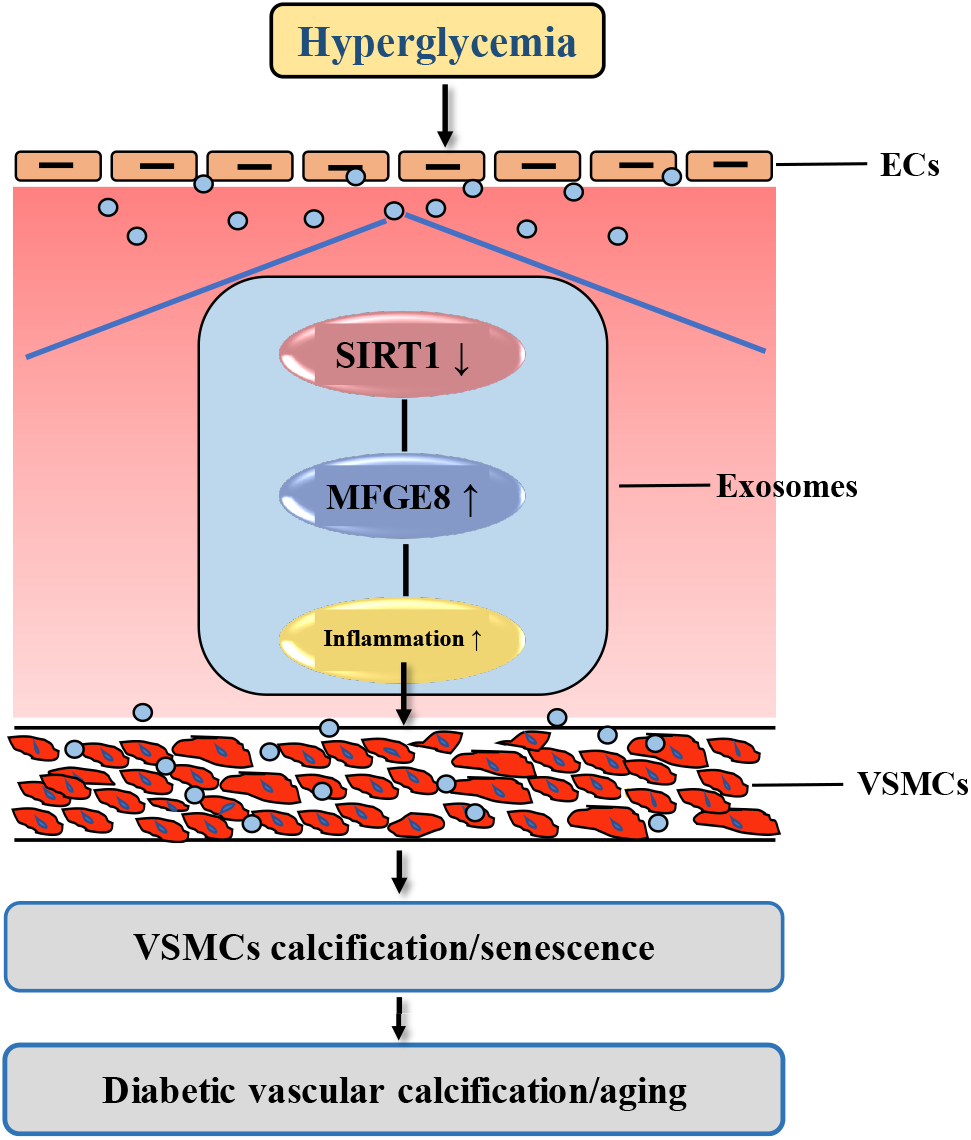
The mechanism diagram of exosomal MFGE8 secreted by ECs in high glucose–induced calcification/senescence of VSMCs. High glucose stimulates the secretion of exosomes by ECs. SIRT1 is deregulated in ECs under hyperglycemia, which in turn regulates the expression of exosomal MFGE8. Then high levels of exosomal MFGE8 are transmitted to VSMCs and promote the calcification/senescence of VSMCs through regulating inflammatory responses, which eventually causing diabetic vascular calcification/aging.

## Materials and Methods

### Cell co-culture

We purchased human VSMCs and HUVEC from the Type Culture Collection of the Chinese Academy of Sciences. VSMCs were maintained in Dulbecco’s Modified Eagle’s Medium (DMEM) with 10% fetal bovine serum (FBS), 1% streptomycin and 1% penicillin. HUVEC were cultivated in F12k medium with 10% FBS, 1% streptomycin and penicillin, and 0.05 mg/ml endothelial cell growth supplement (ECGS). Both HUVEC and VSMCs were cultured at 37 °C, 5% CO2 and saturated humidity. In cell co-cultivation, HUVEC and VSMCs were separated by a membrane with an aperture of 0.4 μm.

### Exosomes isolation

HUVEC were treated with normal glucose (5 mmol/L, NG) or high glucose (30 mmol/L, HG) respectively and inoculated in 55 cm^2^ culture flasks. Exosomes were isolated from the supernatants of HUVEC cultures using differential centrifugation. The supernatants were collected after 48 hours incubation and centrifuged at 2000 *g* for 30 minutes, 10,000 *g* for 45minutes. The supernatants were then passed through a 0.45 μm filter (Millipore) followed by an ultracentrifugation at 100,000 *g* for 70 minutes. The pellets were washed with PBS, ultracentrifuged again at 100,000 *g* for 70 minutes, and resuspended in PBS. All centrifugations were done at 4 °C. Exosomes were utilized for downstream experiments immediately or stored at −80 °C.

### Exosomes identification

The exosomes were resuspended in an equal volume of 10 μL phosphotungstic acid for 1 minute. The morphology and diameter of exosomes were determined by transmission electron microscope (HT-7700, Hitachi) and nanoparticle tracking analyzer (NanoSight NS300, Malvern). Exosomal specific marker proteins CD63 and CD9 were identified by Western blot.

### Calcification assays

Alizarin Red S staining was used to evaluate VSMCs calcification. VSMCs were co-cultured with HUVEC-Exo or HUVEC under NG or HG for 14 days, then fixed with 4% paraformaldehyde for 30 minutes, followed by staining with 1% (pH 4.2) Alizarin Red S for 2 minutes. The mineralized staining were assessed by digital microscope. Besides, the expressions of osteogenic markers ALP and Runx2 by western blot were also used to determine calcification.

### SA-β-gal staining

SA-β-gal staining was performed to determine VSNCs senescence using a SA-β-gal staining kit (Beyotime, Shanghai, China). β-galactosidase fixation solution was used to fix VSMCs for 15 minutes and washed by PBS for 3 times. The percentage of SA-β-Gal positive cells of each sample was determined after staining in β-galactosidase solution at 37°C for more than 12 h.

### Western blot analysis

The cells were lysed with RIPA lysate containing 1% NP-40、0.1% SDS、50 mM DTT, and protease inhibitors including 2ug/ml Aprotinin, 2ug/ml Leupeptin and 1mM PMSF. The homogenate was centrifuged at 9,000 g for 10 minutes and the supernatant was collected. The protein concentration was determined using a NucBuster TM protein Exaction kit (merckmillipore, Cat. No. 71183-3). Then, 30 µg of protein was loaded on a 10% sulphate-polyacrylamide gel electrophoresis (SDS-PAGE) and transferred to a polyvinylidene fluoride (PVDF) membrane. The membranes were incubated with the primary antibody, including ALP (GTX119505, 1:500, Genetex), Runx2 (ab76956, 1:1000, Abcam), CD63 (25682-1-AP, 1:200, Ptgcn), CD9 (20597-1-AP, 1:500, Ptgcn), MFGE8 (29100, 1:1000, SAB), SIRT1 (sc-15404, 1:500, Santa Curz), and β-actin (66009-1-Ig, 1:2000, Ptgcn) at 4 °C overnight after blocking with 5% nonfat milk. After washing the membrane with PBS four times every 5 minutes, the secondary antibody was incubated with the membrane at room temperature for 1 hour. The protein bands were visualized using an enhanced chemiluminescence (ECL) kit and were analyzed by densitometry.

### Quantitative real-time PCR (qRT-PCR) analysis

Total RNA was extracted from VSMCs using Trizol reagent (Invitrogen, 15596-026). To detect MFGE8 and SIRT1 mRNA, cDNA was synthesized from 1 μg of RNA using a RevertAid™ H Minus First Strand cDNA Synthesis Kit (Fermentas, K1631). Then, a 25-μl reverse transcription reaction was performed following the manufacturer’s instructions. SYBR Green PCR Master Mix (ABI 4309155) was used for cDNA amplification in a real-time fluorescence quantitative PCR instrument (ABI, New York, NY, USA). Data were normalized to β-actin values. The PCR primers used in this study were shown in the supplementary Table 1.

### RNA interference

SiRNA-MFGE8 was designed and performed by Gene Pharma Co. Ltd. (Shanghai, China). The sequences of MFGE8 siRNA used are shown in supplementary Table 2. Either MFGE8 siRNA or negative control siRNA was added into the cells and transfected using Lipofectamine 3000 Kit (Thermo Fisher, Waltham, MA, USA). The inhibitory efficiency of siRNA was detected by qRT-PCR.

### Proteomic analysis

The NG-HUVECS-Exo and HG-HUVEC-Exo samples were subjected to iTRAQ-based quantitative proteomic analysis by Jingjie PTM Biolab (Hangzhou, China). High-performance liquid chromatography tandem mass spectrometry (HPLC–MS/MS) was used to detect the different proteomic contents. Differentially expressed proteins were considered significant with a cut-off of absolute fold change ≥1.3 and a corrected P value < 0.05.

### Gene chip analysis

The VSMCs treated with HG-HUVEC-Exo and NG-HUVEC-Exo samples were subjected to gene chip analysis by Kangchen Biolab (Hangzhou, China). R language software was used to read the original data, and Limma package was used for differential gene expression analysis. Differentially expressed genes were considered significant with a corrected P value < 0.05.

### Enzyme linked immunosorbent assay (ELISA)

The levels of pro-inflammatory cytokines IL-1β, IL-6, and IL-8 in cell supernatants were quantified by commercial ELISA kit (EHC002B.48 QuantiCyto, Shenzhen, China), (SEA079Hu, Cloud-Clone CORP, Wuhan, China), and (SEA080Hu, Cloud-Clone CORP, Wuhan, China) respectively, following the manufacturer’s protocol.

### Statistical analysis

All data are presented as means ± SEM and analyzed with Statistical Product and Service Solutions (version 23.0). GraphPad Prism 8.0 is utilized to compare the different groups using one-way ANOVA or t-test. A P-value of < 0.05 was considered statistically significant. All experiments were repeated at least three times.

## Supplementary Material

Supplementary information can be found in the online version of this article.

## Acknowledgements

This work was supported by the National Natural Science Foundation of China (No. 81974223, 82071593, 81770833, and 82101663); the Fundamental Research Funds For the Central Universities of Central South University (NO. 2019zzts354).

## Data Availability

The data used to support the current study are available from the corresponding author upon request.

## Author contributions

Y-SL and J-KZ conceived and designed the experiments; Y-QN carried out most of the experiments and analyzed the data; SL, Y-JW, J-YH, W-LS, YZ, CL, YW, H-HL, ZL provided technical support and discussed the results. Y-QN wrote the manuscript with the help of XL and Q-YX. Y-SL and J-KZ supervised the manuscript. All authors read and approved the final manuscript.

